# An Integrating Sphere Based Method for Quantifying RNA Encapsulation Efficiency in Lipid Nanoparticles without Lysis

**DOI:** 10.64898/2026.05.29.728751

**Authors:** Aida López Espinar, Eric C. Le Ru, Kirsty Smith, Brendan L. Darby

## Abstract

Encapsulation efficiency (EE) is a critical quality attribute of lipid nanoparticle (LNP)-RNA formulations, determining the fraction of active RNA payload available for delivery and influencing therapeutic potency, dose accuracy, and product consistency. Conventional fluorescence-based EE assays, such as RiboGreen (RG), require detergent-mediated LNP lysis, which can introduce systematic bias arising from incomplete particle disruption and additional sample handling steps. Here, we report a lysis-free method for direct EE determination using Scatter-Free Absorption Spectroscopy (SFAS) within an integrating sphere. In this approach, RG fluorescence appears as a quantitative negative absorbance feature in the SFAS spectrum, enabling direct quantification of free RNA from intact LNP suspensions in a single measurement. Applied to SM-102 LNPs loaded with polyA, the method yielded an EE of 97%, in close agreement with conventional plate-reader measurements (98%). This approach provides a practical, formulation-independent, and potentially more robust strategy for standardized routine EE analysis and LNP quality-control workflows.

---

The accurate determination of RNA encapsulation efficiency (EE) in lipid nanoparticle (LNP) formulations remains a significant analytical challenge in the development of RNA-based therapeutics.^1–3^ EE is a critical quality attribute of LNP formulations, directly influencing dose accuracy, protection of RNA from extracellular degradation, intracellular delivery efficiency, and overall product consistency.^1,4^ Reliable quantification of EE is therefore essential throughout both formulation development and quality control.

LNPs have emerged as the leading RNA delivery platform owing to their ability to protect nucleic acids, extend circulation time, and facilitate cellular uptake.^5–7^ Their clinical potential was demonstrated by the rapid global deployment of mRNA vaccines during the COVID-19 pandemic, underscoring their scalability and safety profile,^8,9^ and interest in LNP-based technologies for broader preventative and therapeutic applications has grown considerably since.^5,7,10,11^ LNPs typically comprise four lipid components, an ionizable lipid, a phospholipid, cholesterol, and a PEG-lipid, which self-assemble with RNA within a lipid matrix.^12–14^ Each component serves a defined function: the ionizable lipid mediates endosomal escape, the phospholipid provides structural integrity, cholesterol modulates membrane fluidity, and the PEG-lipid confers colloidal stability.^13,15^ As these systems advance toward clinical use, robust and reliable methods for EE determination become increasingly critical.

The most widely used method for EE determination is the fluorescence-based RiboGreen (RG) assay.^2,3^ The assay relies on the property that RG exhibits negligible fluorescence in solution but undergoes a strong fluorescence enhancement upon binding to RNA, producing a signal proportional to the RNA concentration in the sample.^16^ Operationally, free and total RNA are quantified by measuring RG fluorescence signals obtained before and after detergent-mediated disruption of LNPs, respectively. In intact particles, only non-encapsulated RNA is accessible to the dye, so the fluorescence signal prior to lysis reflects free RNA exclusively. Detergent-mediated lysis then releases encapsulated RNA, enabling measurement of total RNA content.^2,3^ EE is subsequently calculated as the fraction of free RNA in the LNPs suspension relative to the total RNA present.

Despite its widespread adoption, the RG assay is limited by its dependence on complete nanoparticle disruption.^1,16^ The nanoparticle lysis step assumes full release of encapsulated RNA, and it has been well established that this step represents one of the major sources of error for EE quantification.^17–20^ This assumption is not always satisfied, highly stable LNP formulations or those with atypical lipid compositions may resist detergent-mediated disruption, resulting in incomplete RNA release.^16,18^ Direct evidence for this limitation was provided by Schultz et al., who demonstrated that even using different detergents and concentrations, certain formulations were not fully lysed resulting in an underestimation of total mRNA concentrations compared to those obtained by HPLC, and confirming with cryo-TEM imaging that intact LNP structures persisted after exposure to Triton X-100 at both low and high concentrations, confirming the limitations of the detergent based lysis step.^18^ Consistent with this, more studies have shown approximately 20% discrepancy between RG-Triton extraction and LC-MS total mRNA quantification, and questionable accuracy due to the lysis-step.^19,20^ Consequently, post-lysis fluorescence can underestimate total RNA content, leading to an overestimation of EE.^2^ At the same time, the assay retains a useful and often underexploited feature. In the absence of nanoparticle lysis, RG selectively reports RNA that is accessible in solution. Because the dye cannot cross intact lipid membranes, fluorescence measured in intact LNP suspensions corresponds directly to non-encapsulated RNA.^16^

To address the limitations of lysis-dependent approaches, methods that enable direct RNA quantification in intact nanoparticle suspensions are required.^1^ Scatter-free absorption spectroscopy (SFAS) is a UV/Visible (UV/Vis) technique developed for this purpose.^21,22^ Conventional UV/Vis measurements are confounded by strong light scattering from nanoparticles, which distorts the baseline and interferes with absorbance-based RNA quantification at 260 nm.^21,23,24^ SFAS overcomes this limitation through the use of an integrating sphere that collects transmitted and scattered light, enabling measurement of total light intensity independent of detection geometry.^21,24,25^ This configuration removes scattering artefacts on UV absorbance, yielding a “true” absorbance spectrum of the sample. Because of the complex pathlengths of rays inside the integrating sphere, this measurement must be calibrated and corrected to yield the same absorption coefficient as would be observed in standard UV/Vis over a 1 cm pathlength.^24,25^ This scatter-free-absorbance (SFA) spectrum can then be used, via the Beer-Lambert Law, to directly quantify total RNA in intact LNP suspensions without needing to lyse the particles.^21,22^ As a result, SFAS provides robust measurements of total RNA concentrations in LNPs that do not require nanoparticle disruption. SFA has been validated across a range of nanoparticle formulations and payload types, regardless of particle size or lipid composition.^21,22^ However, SFAS as it is currently implemented only reports total RNA concentration and does not distinguish between encapsulated and non-encapsulated RNA populations.

In this work, we present a combined analytical strategy that integrates SFAS-based RNA quantification, with fluorescent dye detection to enable lysis-independent determination of RNA encapsulation efficiency. The method relies on two orthogonal measurements performed directly on intact LNP suspensions within the same integrating sphere configuration: (i) total RNA is quantified by standard SFAS, which corrects for light scattering contributions and isolates the RNA absorbance signal at 260 nm, and (ii) we demonstrate that the SFAS signal can also be used to quantify fluorescence and therefore free RNA when RG is added to intact LNP suspension. The dye selectively binds to non-encapsulated RNA, and the resulting fluorescence emission is detected within the integrating sphere measurement, appearing as a characteristic negative feature in the absorbance spectrum. This enables free RNA to be quantified from the same SFAS instrument without a separate fluorimetric measurement.

EE is then calculated directly from these two orthogonal signals obtained within a single optical platform. By eliminating the need for nanoparticle lysis and exploiting fluorescence detection within an absorption-based integrating sphere measurement, this approach removes a major source of systematic error in conventional EE assays, avoids dependence on formulation-specific lysis efficiency, and provides a more robust and broadly applicable strategy for characterizing RNA-loaded LNP systems. Although the approach is applicable to RNA broadly, the experiments presented here were performed using polyadenylic acid (polyA) as a model of RNA, owing to its commercial availability and its well-characterized optical properties.^26^

## Detection of RiboGreen Fluorescence by Scatter-Free Absorption Spectroscopy

The RG assay is based on fluorescence emission upon binding of the RG dye to RNA (or polyA) (Figure 1A), where the fluorescence signal intensity is proportional to the amount of accessible RNA that binds to the RG (Figure 1B).^16^ In conventional plate-reader based measurements, the sample is excited at a defined wavelength, typically at the absorption maximum of the dye. Fluorescence is detected at a wavelength higher than the excitation wavelength. In the case of RG dye, the excitation and detection wavelengths are typically 480 nm and 520 nm respectively (Figure 1A),^16^ corresponding to where the absorption and fluorescence of the dye are the highest.

**Figure 1.**
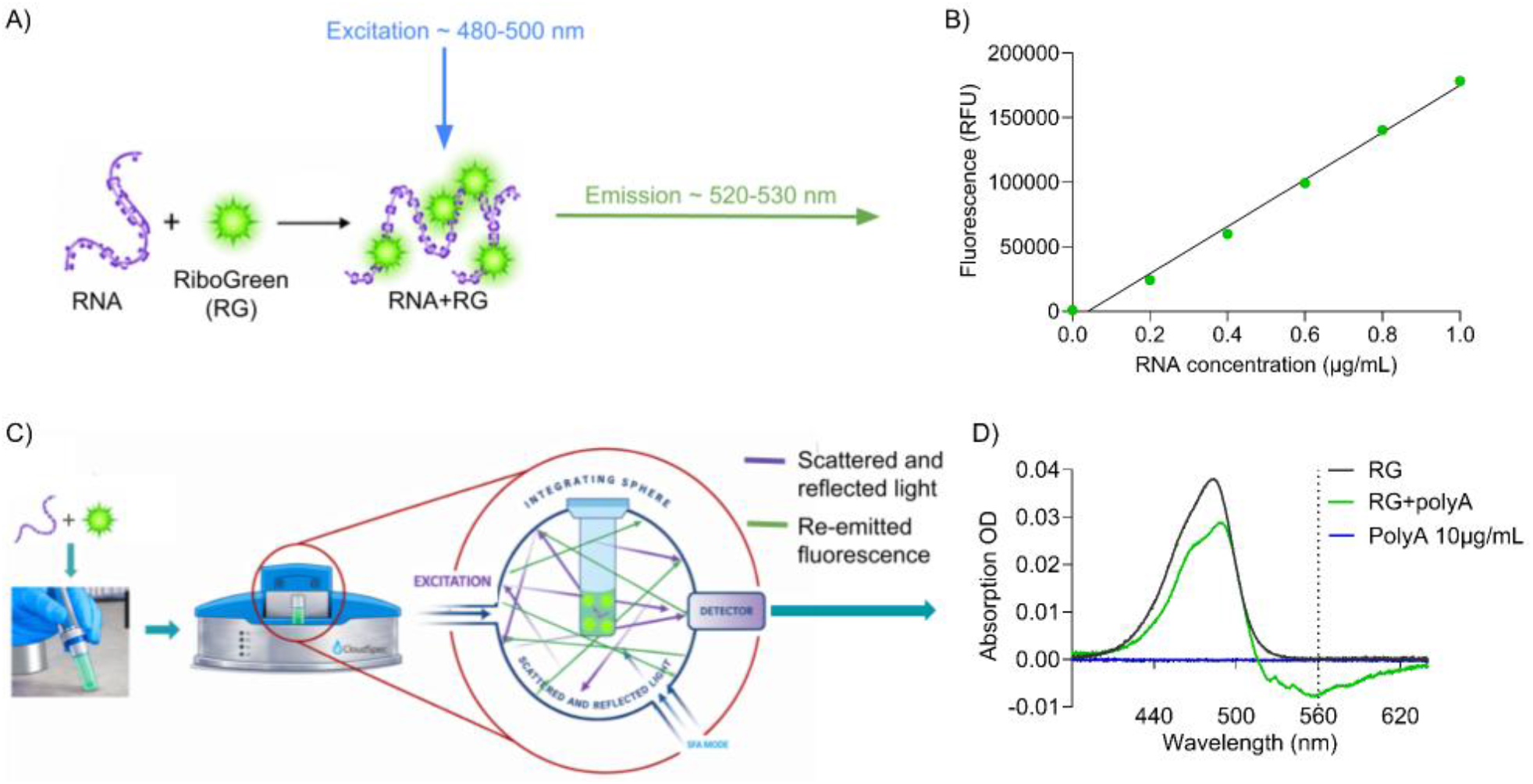
A) RNA and RiboGreen (RG) forming a fluorescent complex with defined plate reader fluorescence excitation (∼480-500 nm) and emission (∼520-530 nm) wavelengths. B) Plate reader measured RG Relative Fluorescence Units (RFU) increasing linearly with RNA concentration. C) Schematic of an SFAS measurement showing the integrating sphere that collects transmitted, scattered, and emitted light. D) SFAS spectra for RG-polyA complex (green line), RG (black line) and polyA (10 μg/mL) (blue line).

We investigated how this fluorescence signal is captured using SFAS, which relies on a fundamentally different detection principle. In SFAS, the sample is placed within an integrating sphere that collects all optical contributions, including transmitted, scattered, and emitted light (Figure 1C). It is excited by broadband excitation, typically from UV to near infrared. Upon illumination, the excitation light interacts with the sample, where it can be scattered, absorbed and, in the presence of RG-bound to RNA (or polyA) re-emitted as fluorescence (Figure 1C). The integrating sphere redistributes this light through multiple diffuse reflections, enabling detection of the total light intensity independent of direction.^21^

Within this framework, fluorescence emission from RG-bound to RNA contributes directly to the detected SFAS signal. The emitted light increases the overall intensity reaching the detector at the fluorescence wavelengths, which in SFAS appears as a decrease in absorbance. In spectral regions where fluorescence does not overlap with absorption, this will be observed as an apparent negative absorbance feature directly related to the fluorescence signal. In the presence of polyA, a clear negative absorbance signal is therefore observed in the spectra, peaking around 560 nm (Figure 1D, green line). Note that the peak here is shifted compared to a conventional measurement (where it is usually seen at 520 nm) for at least two reasons. Firstly, the excitation is now much broader, including wavelengths overlapping with the normal fluorescence spectrum. Secondly, RG absorbance extends up to ∼550 nm and will therefore partly cancel the negative fluorescence spectrum in this region. Similar wavelength-dependent changes in fluorescence lineshapes have also been observed previously in integrating-sphere measurements, where multiple-pass excitation and omnidirectional collection modify the detected emission profile compared to conventional single-pass fluorimetry.^27^

Control experiments confirmed that the SFA fluorescence feature is only observed when both polyA and RG are present and is absent in samples containing either RG alone (Figures 1D, black line) or polyA (10 μg/mL) alone (Figure 1D, blue line), demonstrating that it originates specifically from RG-RNA interactions.

### Theoretical Description of Fluorescence in SFAS

In the presence of fluorescence, the SFAS spectrum is a complex mixture of the absorbance spectrum (positive) and the negative of the fluorescence spectrum. Deconvolving these two contributions can be difficult. However, in regions where sample absorbance is negligible, such as at wavelengths beyond the absorption peak (***λ***>550nm in the case of RG), the increase in intensity is directly proportional to the fluorescence spectrum as explained below. The detected intensity for the reference (here TE 1X buffer) can be written as *I*_*ref*_ (*λ*), and for the sample (here RG in TE 1X) as *I*_*ref*_ (*λ*) + *α I*_*fluo*_(*λ*). The SFAS spectrum is reported as optical density (OD) corrected for effective pathlength, which introduces a more complex relationship to the fluorescence intensity. The raw OD is defined as:

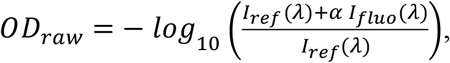

and the corrected OD is:

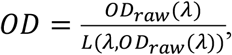

where the effective pathlength *L* depends on wavelength and measured OD.^24,25^

For low fluorescence signals compared to the excitation source, i.e. when *αI*_*fluo*_(*λ*) ≪ *I*_*ref*_ (*λ*), these expressions can be simplified to recover a simple proportionality relation between SFAS signal and fluorescence intensity:

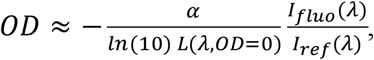

Which shows that |OD| is proportional the fluorescence signal. This approximation is valid for *αI*_*fluo*_(*λ*)/*I*_*ref*_(*λ*) ≤ 0.23, with an error under 10%. For our instrument, the effective pathlength at zero absorbance is *L*(*λ, OD* = 0) ≈ 4 *cm*, so the linear approximation is valid for |*OD*| ≤0.025. As will be shown, the RG fluorescence measurements at concentrations relevant to the assay fall within this range. While these ODs may appear quite small, the integrating sphere enables their measurement with high precision, owing to its averaging effect on optical imperfections and the pathlength enhancement it provides. This suggests that the SFAS signal at 560 nm can be used to quantify the RG fluorescence in the same way as with a plate reader excited at 480 nm and read at 520 nm, with the signal proportional to fluorescence intensity in both cases.

### PolyA quantification using RiboGreen with SFAS

To further demonstrate this, we evaluated the relationship between the RG SFAS fluorescence (OD at 560nm) and polyA concentration following the recommended protocol for the high-range assay: RG reagent diluted 400X in TE 1X buffer. The reported linear range is then 0.1 μg/mL<cpolyA<1 μg/mL.^16^ As shown in Figure 2A, the magnitude of OD560 increases progressively with increasing polyA concentration, demonstrating a clear and systematic concentration-dependent response. This indicates that the fluorescence contribution detected by SFAS can be used to quantitatively measure accessible RNA. The precision of the OD560 measurement, expressed as the standard deviation over three replicate measurements, was under 2% across all polyA concentrations tested. Importantly, the signal at 560 nm shows a strong linear correlation with polyA concentration across the tested range (Figure 2B), which is a prerequisite for its use as a quantitative analytical tool. This linearity is consistent with the expected behavior of the RG assay, where fluorescence intensity scales proportionally with RNA concentration within a defined working range.^16^ This is also supported by the agreement between the increase in fluorescence signal observed with increasing polyA concentration for the same samples measured using a regular plate reader (Figure 2C), which confirms that the two approaches are equivalent. The direct linear correlation observed between SFAS fluorescence intensity and plate reader fluorescence (Figure 2D) further confirms quantitative equivalence between the two approaches across the concentration range tested. This equivalence is consistent with previous demonstrations that integrating sphere detectors can quantitatively capture fluorescence emission from solution-phase samples with accuracy comparable to conventional fluorimeters.^27^

**Figure 2.**
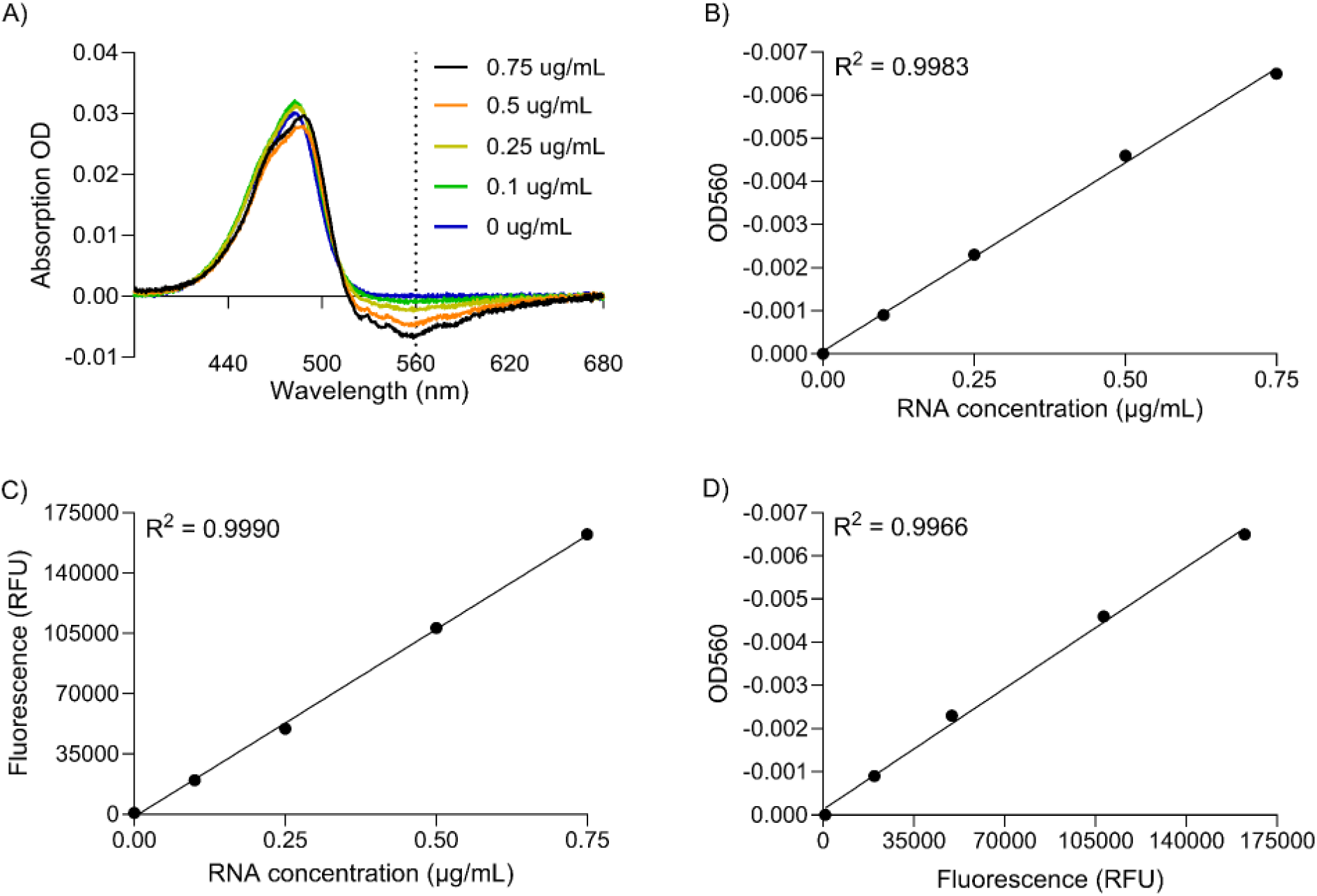
A) SFAS spectra of RG at increasing polyA concentrations. B) Linear correlation between the SFAS signal at 560 nm and polyA concentration. C) Linear correlation between Relative Fluorescence Units (RFU) measured by plate reader and polyA concentration. D) Linear correlation between SFAS signal (OD at 560 nm) and plate reader RFU.

These results establish that OD560 in SFAS is a reliable and quantitative reporter of RNA concentration using the RG assay. The linearity and agreement with conventional fluorimetry validate its use as an orthogonal detection channel within the SFAS measurement, enabling RNA quantification without a separate fluorescence instrument.

### Quantifying Free polyA in LNPs

In LNP formulations, RNA exists in two distinct populations: encapsulated RNA, which is protected within the nanoparticle interior and shielded from the external aqueous environment, and non-encapsulated (free) RNA, which remains accessible in the surrounding solution.^1,28^ This distinction is analytically critical because only encapsulated RNA is protected from extracellular nuclease degradation and available for intracellular delivery, while free RNA contributes to the total nucleic acid content of the formulation without contributing to therapeutic efficacy.^29^

The RG assay is commonly used in conjunction with a plate reader to quantify free RNA in LNP solutions. As the RG can only bind to the free RNA in solution, not the LNP-encapsulated RNA, the method provides a direct measure of the free RNA concentration.^2,16^ Here, we demonstrate that the same approach can be followed with SFAS.

One complication that is often overlooked is that the RG assay is strongly buffer-dependent.^1,3,17^ Since LNPs are often suspended in non-TE buffer such as PBS 1X or sucrose-containing buffer, it is not practical to prepare a RG+LNP in pure TE 1X (this would require dialysis of the LNPs in this buffer). It is often assumed that a sufficiently high dilution in TE 1X will be equivalent to pure TE 1X, but as shown in Figure 3A, this is not the case. Even a small amount of PBS results in (i) a decrease in the fluorescence emission, and (ii) a reduction in the linear range, here down to cRNA<0.5 μg/mL).^30^ This behavior is consistent with the known sensitivity of RG fluorescence to ionic strength, where increased salt concentration can screen electrostatic interactions between the positively charged dye and the negatively charged RNA backbone, thereby reducing dye binding affinity and fluorescence quantum yield.^16,30^ When not carefully accounted for, such effects are likely the source of some of the irreproducibility problems often reported with the RG assay.^3,17^

**Figure 3.**
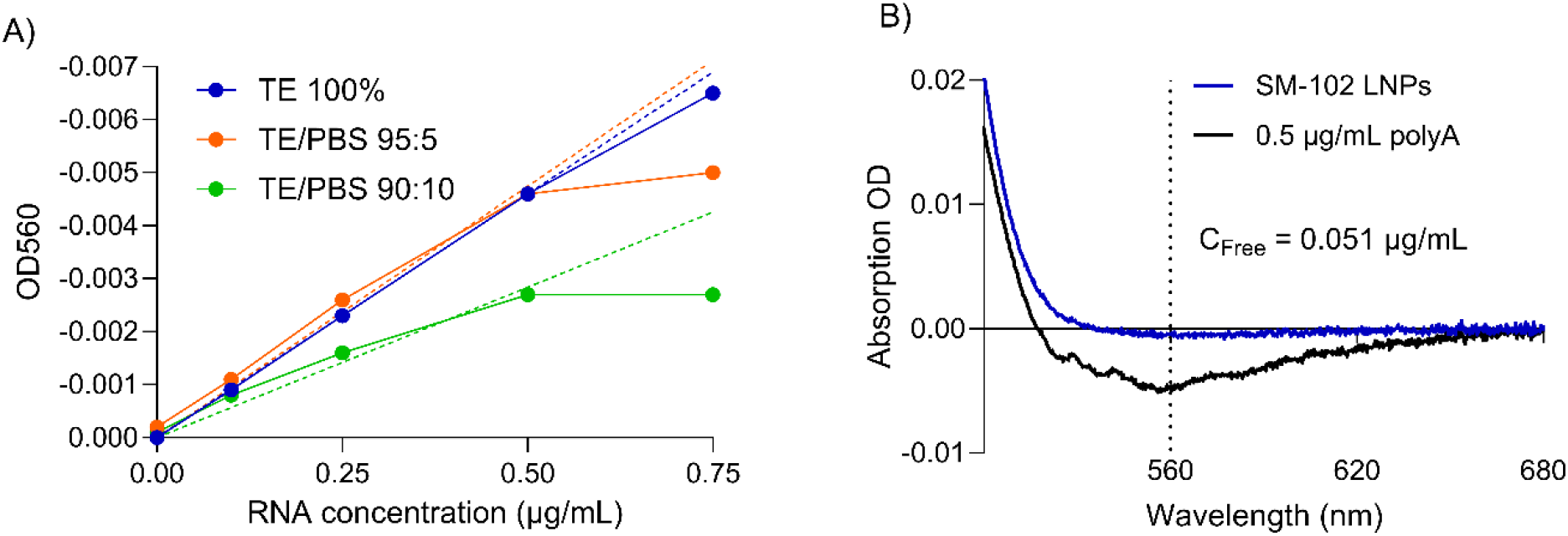
A) RG fluorescence as quantified by OD560 in SFAS as a function of RNA concentration in different buffers: pure TE, 95:5 TE/PBS, and 90:10 TE/PBS. The dashed lines show linear fit through the origin based on concentrations up to 0.5 μg/mL to highlight the deviation from the linear range at 0.75 μg/mL. B) SFAS spectra of SM-102 LNPs (blue line) and reference polyA (0.5 µg/mL) (black line).

In the following, we used the 95:5 TE/PBS ratio for all measurements (including calibration curves) as it still provides a fluorescence similar to pure TE 1X (Figure 3A). This will require a 20X dilution of the LNP with sample preparation as follows:

1045 μL of 380X RG reagent in TE 1X + 55 μL of LNP in buffer (PBS 1X here)

This provides a 400X RG reagent final concentration as required in 95:5 TE/PBS. There is enough sample for the 1 mL volume required for the SFAS measurement with some margin for more accurate pipetting.

Based on the observed reduced linear range (Figure 3A), we also prepared a reference sample of known polyA concentration at 0.5 μg/mL, also in 95:5 TE/PBS as follows:

1045 μL of 380X RG reagent in TE + 55 μL of 10 μg/mL polyA

Note that it is recommended to use the same RNA for the calibration sample as for the LNP since the RG fluorescence is known to vary with the RNA type.^2,17^

We applied the proposed approach to SM-102 LNPs encapsulating polyA (Figure S1, Supporting Information). SM-102-based LNPs are well-characterized model systems widely used in RNA delivery research and form the basis of clinically approved mRNA vaccines, making them a relevant and practically meaningful benchmark for this analysis.^8,9,31^ The SFA fluorescence of the RG+LNP and RG+polyA samples are shown in Figure 3B. Based on the linear dependence of SFA at OD560 with free polyA concentration, we derive the free polyA concentration in the original LNP solution, accounting for any dilution as:

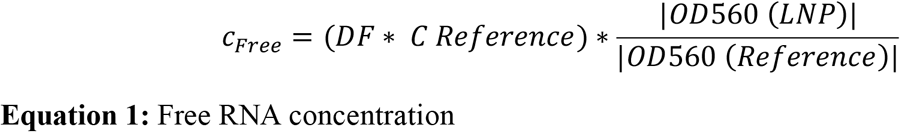

In the example of Figure 3B, we have C_Reference_ = 0.5 μg/mL and DF = 50 (arising from a 2.5-fold pre-dilution of the LNPs to get 55 µL at a target concentration of 40 µg/mL, combined with a 20-fold dilution upon addition of the RG reagent), giving a C_Free_ of 0.051 µg/mL in the measured sample equivalent to C_Free_=2.54 µg/mL in the original LNPs stock (Figure 3B).

Note that preparing the reference in the same buffer as the LNP sample ensures that any buffer-dependent modification modulation of the RG fluorescence response, such as the signal reduction and narrowing of the linear range observed in the presence of PBS, affects both the reference and the sample equally, and is therefore cancelled in the ratiometric calculation.^17^ This matrix-matched referencing strategy directly addresses the buffer sensitivity identified earlier and is essential for obtaining accurate free RNA concentrations independently of the specific buffer composition of the LNP formulation. This is true whether the fluorescence is measured with SFAS or a plate reader. Our choice of a 95:5 TE/PBS ratio is a compromise. A higher dilution factor, such as 98:2 may not be practical as it requires a higher LNP concentration to start with. But a lower dilution such as 90:10 results in a decrease in fluorescence (Figure 3A). The 95:5 ratio provides the same fluorescence intensity as pure TE 1X (Figure 3A). The only trade-off is that the linear range is reduced to cRNA<0.5 µg/mL (Figure 3A). But with the pathlength enhancement in the integrating sphere, the lower detection limit is improved, so the working range is 0.05 µg/mL<C_Free_<0.5 µg/mL, or equivalently 1 µg/mL<C_Free_<10 µg/mL before the 20x dilution.

### Determining Encapsulation Efficiency using SFAS

The EE quantifies the proportion of RNA that is successfully incorporated into the nanoparticle structure and is therefore a primary quality attribute of any LNP formulation, directly informing dose accuracy, therapeutic potency, and formulation quality.^1,2,32,33^ EE is defined according to Equation 2:

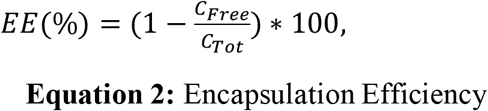

where C_Free_ is the concentration of non-encapsulated accessible RNA and C_Tot_ is the total RNA concentration. The accurate EE determination therefore requires independent, reliable quantification of both quantities. In the SFAS-based approach presented here, C_Free_ is determined from the SFAS at 560 nm arising from RG-RNA as just explained. C_Tot_ is measured independently from the SFAS RNA absorbance signal at 260 nm on intact LNP suspension, without detergent-mediated particle.^21,22^ This preserves sample integrity and eliminates the artefacts associated with incomplete or heterogeneous LNP lysis.

An important point is that different sample dilutions are typically used for the determination of free RNA concentration (C_Free_) and total RNA concentration (C_Tot_). For the determination of C_Free_ (RG + LNPs), samples were prepared to achieve a targeted concentration of C_Tot_ around 2 μg/mL (i.e. 40 μg/mL before the 20X dilution in RG/TE needed for the sample preparation). Given the linear detection range of the assay (0.05 <C_Free_ <0.5 µg/mL), this concentration range enables accurate determination of EE between approximately 75% and 97.5%, which covers the range typically relevant for LNP formulations. This range can be adjusted depending on the application requirements. For example, a target concentration of C_Tot_ = 1 µg/mL after dilution (20 µg/mL prior to dilution) would allow quantification within an EE range of approximately 50% to 95%.

As pointed out before, the determination of C_Tot_ by SFAS is an independent measurement, where the LNP suspension is diluted to a concentration appropriate for SFAS absorbance measurement at 260 nm, corresponding to an RNA concentration in the range 2 to 20 µg/mL. In the representative example shown in Figure 4A, the SM-102 LNP stock (nominal concentration ∼100 µg/mL was diluted 10-fold). The SFAS measurement gives a measured RNA concentration of 8.36 µg/mL, and therefore a C_Tot_ of 83.6 ± 0 µg/mL in the original stock (Figure S2, Supporting Information). The small reduction relative to the nominal initial formulation concentration is due to dilution during dialysis. For comparison, C_Tot_ quantification of the same sample using the conventional RG assay with lysis measured in a plate reader (C_Tot, PR_) yielded 66.7 ± 2.5 µg/mL, approximately 20% lower than the SFAS-derived value and with greater measurement variability (Figure S2, Supporting Information). This systematic underestimation is consistent with incomplete LNP lysis, a recognized limitation of detergent-based disruption protocols,^1,18^ and is avoided when total RNA is quantified directly by absorbance on intact particles.

**Figure 4.**
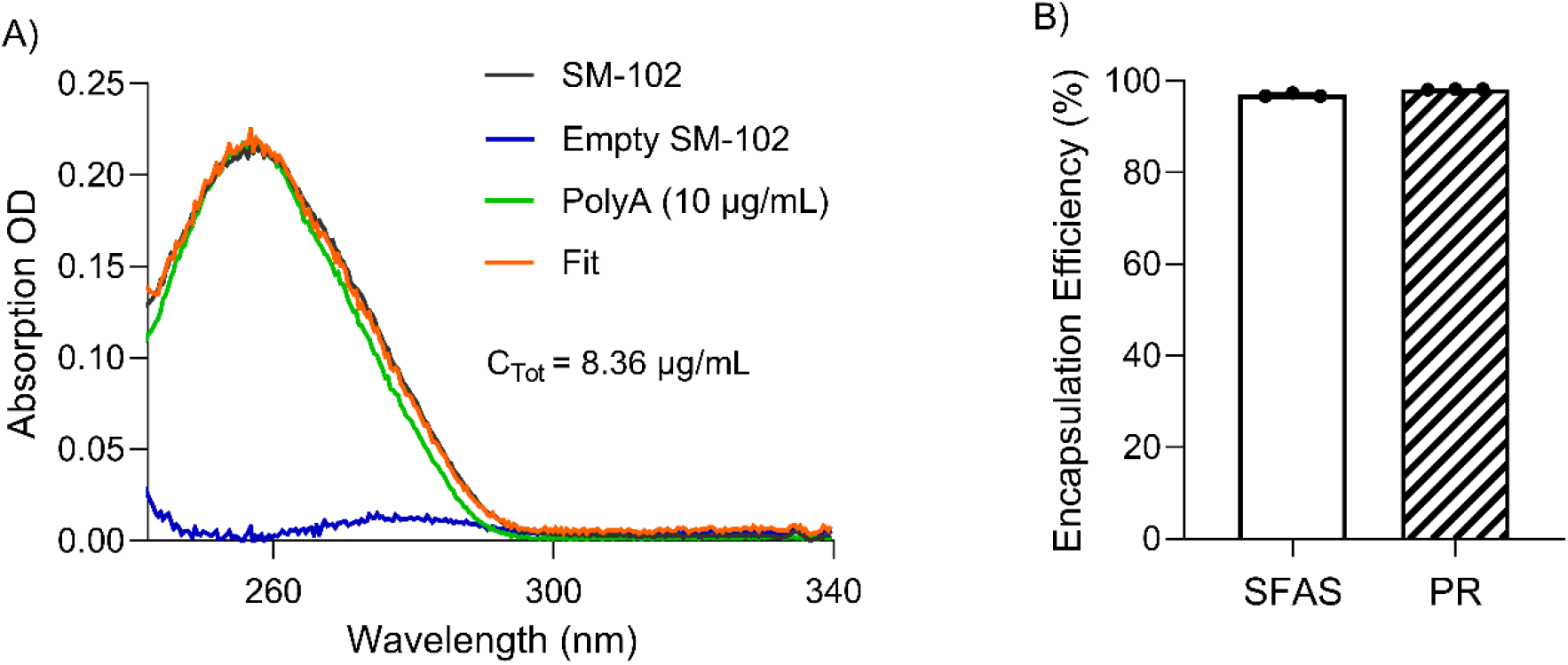
A) SFAS of polyA-loaded SM-102 LNPs (black line). The total RNA concentration is determined by performing a deconvolution of this spectrum as a weighted sum of the SFAS spectra of empty SM-102 LNPs (contribution shown as blue line) and free polyA (10 μg/mL) (green line). The deconvolution fit is shown as orange line. B) Encapsulation efficiency determined by SFAS and PR.

EE is then calculated after accounting for the respective dilution factors. Using C_Free_ and C_Tot_ both determined by SFAS, an EE of 97% was obtained (Figure 4B). As an independent check, C_Free_ was also determined using the conventional plate reader RG assay on intact, non-lysed LNP suspensions, yielding EE_PR_= 98% (Figure 4B), in close agreement with the all-SFAS result.

## Summary and Conclusions

The EE values derived from SFAS (97%) and the conventional plate-reader RG assay (98%) are in close agreement, validating the proposed approach. A key novelty of this work is that RG fluorescence, conventionally measured using dedicated fluorescence instrumentation, is detected directly within the SFAS measurement as a quantitative negative absorbance feature. This enables both free RNA and total RNA to be determined from the same integrating sphere set-up or instrument. In standard RG-based EE determination, lysis with detergents such as Triton X-100 assumes complete membrane disruption, an assumption that has been shown to fail for stable or atypically composed LNP formulations, leading to systematic underestimation of total RNA and overestimation of EE.^1,18^ By eliminating the lysis step entirely, the SFAS-based approach removes one of the principal sources of error in conventional EE assays and is therefore inherently less sensitive to formulation-dependent variability, making it more robust across diverse LNP compositions in both development and quality-control settings.^21,22^ Compared with the plate-reader assay, SFAS has a higher limit of detection for free RNA (0.05 µg/mL versus ∼1 ng/mL for fluorescence plate readers). Nevertheless, for most LNP formulations, where free RNA concentrations are typically in the µg/mL range, this represents an acceptable trade-off.

An important and often overlooked practical finding is the strong dependence of the RG signal on buffer composition. Even modest amounts of PBS in the assay mixture reduce fluorescence intensity and narrow the linear working range. This behavior is consistent with the known sensitivity of RG fluorescence to ionic strength, where increased salt concentration screens electrostatic interactions between the positively charged dye and the negatively charged RNA backbone, thereby reducing dye binding affinity and fluorescence quantum yield.^16^ Such effects likely contribute to the irreproducibility frequently reported for the RG assay.^1,17^ Consistent with this, Parot et al. identified sample preparation-induced variability as a principal source of bias in fluorescence-based mRNA quantification for LNP formulations, further highlighting the importance of matrix-matched calibration strategies such as those employed here.^19^ Preparing calibration standards in the same buffer composition as the LNP samples directly compensates for these effects through ratiometric cancellation and should therefore be considered best practice regardless of the detection platform used.^13^

In conclusion, we have proposed and demonstrated a lysis-free method for determining RNA encapsulation efficiency in LNP formulations using scatter-free absorption spectroscopy. By combining total RNA quantification with free RNA quantification via directly detected RG fluorescence within the same integrating-sphere measurement, EE can be obtained directly from intact nanoparticle suspensions with precision and accuracy comparable to conventional methods, without requiring nanoparticle disruption. This approach therefore offers a practical and potentially more robust alternative for routine EE determination in LNP formulation development and quality-control workflows.

## ASSOCIATED CONTENT

Supporting Information including:

Figure S1. Size (nm) and Polydispersity Index (PDI) of SM-102 LNPs encapsulating polyA.

Figure S2. SM-102 LNPs total RNA concentration determined by Scatter-Free Absorption

Spectroscopy (SFAS) and Plate Reader (PR) with the standard RG assay.

Materials and methods

Materials

Formulation of SM-102 nanoparticles

Free RNA Quantification with SFAS

Total RNA concentration Quantification with SFAS

Free RNA quantification using RG assay in the plate reader

## AUTHOR INFORMATION

### Author Contributions

The manuscript was written through contributions of all authors. All authors have given approval to the final version of the manuscript.

## ABBREVIATIONS

EE: Encapsulation efficiency
LNP: Lipid Nanoparticle
SFAS: Scatter-Free Absorption Spectroscopy
C_Free_: Free RNA Concentration
C_Tot_: Total RNA Concentration

## TOC

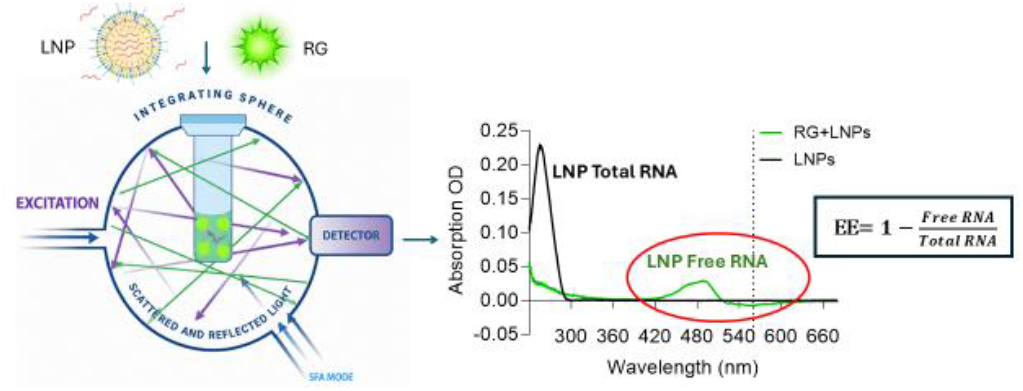

