## Supporting Information for "An Integrating Sphere Based Method for Quantifying RNA Encapsulation Efficiency in Lipid Nanoparticles without Lysis"

### Table of Contents


- **Figure S1.** Size (nm) and Polydispersity Index (PDI) of SM-102 LNPs encapsulating polyA.
- **Figure S2.** SM-102 LNPs total RNA concentration determined by Scatter-Free Absorption Spectroscopy (SFAS) and Plate Reader (PR) with the standard RG assay.
- **Materials and methods**
  - Materials
  - Formulation of SM-102 nanoparticles
  - Free RNA Quantification with SFAS
  - Total RNA concentration Quantification with SFAS
  - Free RNA quantification using RG assay in the plate reader

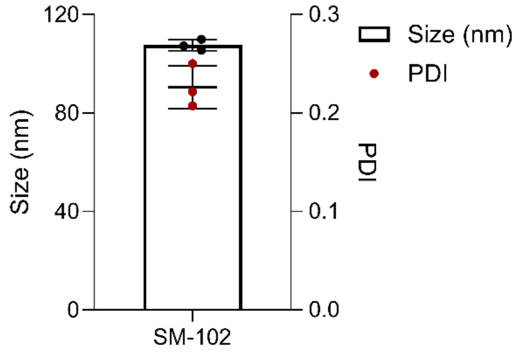


**Figure S1.** Size (nm) and Polydispersity Index (PDI) of SM-102 LNPs encapsulating polyA.

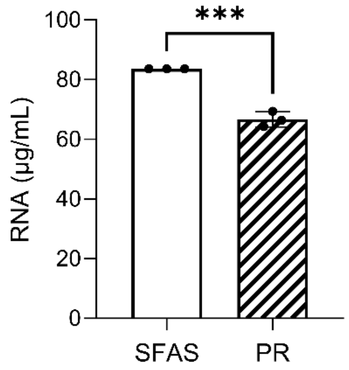


**Figure S2.** SM-102 LNPs total RNA concentration determined by Scatter-Free Absorption Spectroscopy (SFAS) and Plate Reader (PR) with the standard RG assay.

### Materials and methods

Materials

The lipids 1,2-dioctadecanoylsn-glycero-3-phosphocholine (DSPC) and 1,2-Dimyristoyl-rac-glycero-3-methoxypolyethylene glycol-2000 (DMG-$PEG_{2000}$) were purchased from Avanti Research. Cholesterol was purchased from Merck Ireland. The lipid SM-102 was purchased from AVT Pharmaceutical Shanghai. The polyA was purchased from Merck Ireland, manufactured by Roche. The Quant-it™ RiboGreen Reagent, Triton X-100 98% for molecular biology, DNAse, RNAse and Protease free, Citrate 0.5M (pH 3), PBS 10X and Invitrogen™ UltraPure™

DNase/RNase-Free Distilled Water were purchased from Thermo Fisher Scientific Ireland. 20X TE Buffer (pH 7.5) was purchased from Promega Ireland. Ethanol absolute ≥ 99.8% analytical reagent grade was purchased from Merck Ireland.

Formulation of SM-102 nanoparticles

SM-102 LNPs were formulated by stir-bar mixing of ethanol and aqueous phases at a volumetric ratio of 1:3. The mixture was stirred at 1000 rpm for 30 s. The ethanol phase contained SM-102, DSPC, cholesterol, and DMG-PEG2000 at a molar ratio of 50:10:38.5:1.5, respectively. Formulations were prepared at an N/P ratio of 6:1. The aqueous phase consisted of polyA dissolved in a 10 mM citrate buffer (pH 3.2).

Following formulation, SM-102 LNPs were dialyzed in 1X PBS for 4 h using Slide-A-Lyzer Mini Dialysis Devices (20 kDa MWCO, Fisher Scientific Ireland), with the dialysis buffer replaced after 2 h. After dialysis, particle size and polydispersity index (PDI) were characterized by dynamic light scattering (DLS) using a Zetasizer Ultra (Malvern Instrument, Worcestershire, UK) at 25 °C. To determine the size and PDI, samples were prepared by diluting 50 µL of LNPs in 950 µL of PBS 1X. Empty SM-102 LNPs were prepared and characterized in the same way, replacing the polyA by RNase-free water.

Free RNA Quantification with SFAS

Free RNA concentration was quantified from the SFA fluorescence signal at 560 nm (680 nm used as background correction) using a CloudSpec instrument (Marama Laboratories, Ireland). Concentration was determined by a ratiometric approach relative to a matrix-matched reference of known RNA concentration (0.5 µg/mL), according to Equation 1, where DF is the dilution factor of the LNP stock to the 2 µg/mL working concentration and CReference is the reference standard concentration.

A RiboGreen working stock was prepared by diluting the RG reagent 380-fold in TE 1X buffer. The reference sample was prepared by mixing 1045 µL of RG working stock with 55 µL of 10 µg/mL RNA in PBS 1X, yielding a final RNA concentration of 0.5 µg/mL and a 400-fold RG dilution in 95:5 TE/PBS. For SM-102 LNP samples, the formulation was first diluted to ~40 µg/mL RNA in PBS 1X, 55 µL of this was then combined with 1045 µL of RG working stock, giving a final 50-fold dilution of the LNP stock in 95:5 TE/PBS. All samples were prepared to a total volume of 1.1 mL and protected from light throughout. SFAS measurements were performed using a 1 $cm^2$ quartz cuvette with 95:5 TE/PBS as blank and a fresh baseline acquired before each measurement.

Total RNA concentration Quantification with SFAS

SM-102 LNPs were diluted to 10 µg/mL RNA in PBS 1X for SFAS measurement. Empty LNPs were diluted identically. Measurements were performed using a CloudSpec instrument with a 1 $cm^2$ quartz cuvette (1 mL volume). The integrating sphere automatically corrected for pathlength modification effects to report the absorption coefficient as equivalent optical density over 1 cm.[1,2] Extinction spectra were recorded simultaneously in standard transmission configuration; scattering spectra were obtained by subtracting absorption from extinction. The blank was the corresponding buffer for each sample. The RNA-loaded spectrum was fitted to a weighted sum of pure RNA and empty LNP reference spectra; total RNA concentration was determined from the RNA weight coefficient and the known reference concentration, with the 10-fold dilution factor applied to recover the stock concentration.

Free RNA quantification using RG assay in the plate reader

The Quant-iT RiboGreen assay (ThermoFisher) was performed according to the manufacturer's protocol. The assay was conducted in a flat-bottom black 96-well plate (Thermo Scientific™ Sterilin™). A standard curve was prepared by diluting RNA in TE 1X buffer to final concentrations ranging from 0 to 200 ng/100µL. The LNPs samples were diluted to a final volume of 100 µL in TE 1X. Then, 100 µL of RiboGreen (diluted 1:200 in TE 1X) was added to both the samples and the standard curve. The plate was incubated in the dark for 5 min, shaking at 100 rpm using an orbital shaker (Benchmark Scientific). After incubation, the fluorescence intensity of non-encapsulated RNA was measured using a VANTAstar Microplate reader (BMG LABTECH) at an excitation wavelength of 485 (±20) nm and an emission wavelength of 535 (±25) nm. Next, 22 µL of 1% Triton X-100 was added to each sample, and the plate was incubated for 3 min, shaking at 100 rpm in the plate reader. Following incubation, the fluorescence intensity of total RNA was measured under the same conditions.
